# Topological defects and coherent myocardial chirality shape torsional heart contraction

**DOI:** 10.64898/2026.04.13.718243

**Authors:** Naofumi Kawahira, Takaki Yamamoto, Takumi Washio, Yoshiro Nakajima, Kenta Yashiro, Vincent Xu, Kyogo Kawaguchi, Atsushi Nakano

**Author notes:** These authors contributed equally to this work (Naofumi Kawahira, Takaki Yamamoto).

## Abstract

The efficient pumping of the mammalian heart relies on torsional contractile motion generated by its highly ordered three-dimensional (3D) architecture of myocardial fibres. However, the topological principles governing how its complex geometry translates into contractile mechanics remain elusive. Here, we show that the mammalian heart forms a chiral nematic field, a biological analogue to 3D liquid crystals, whose topological organisation underlies its function. Analysis of 3D imaging data revealed disclination lines, continuous assemblies of topological defects characteristic of nematic systems, within the compact myocardium. Finite-element simulations reveal that these defects are not mere structural irregularities but can locally modulate contractile behaviour and reduce mechanical work. In heterotaxy hearts with reversed global anatomy (situs inversus), myocardial fibres retain a predominantly counter-clockwise twist, similar to that of the normal heart, but with a small clockwise component near the base. This decoupling of tissue-level chirality from systemic left-right patterning suggests that cardiac twist is an intrinsic property of the myocardial fibre. Mechanical simulations of situs inversus heart demonstrate that the coherence of transmural chirality, rather than its specific orientation, is critical for contractile efficiency. Together, these findings establish the heart as a topological material and reveal how organised chiral fields generate robust organ-level mechanical function.

---

The heart is a highly organized muscular organ whose efficient pumping relies on the three-dimensional (3D) architecture of its contractile fibres. Cardiomyocytes generate force by shortening along their long axis, and the collective orientation of these elongated cells forms a coherent global field that twists across the myocardium. This transmural twist, forming a counter-clockwise (CCW) chiral configuration in normal human hearts, has been implicated in cardiac contractile performance [1–5]. However, most work to date has focused on anatomical description rather than the physical rule underlying myocardial fibre architecture [6, 7]. Debates remain as to the existence of discrete myocardial band and mechanistic implication of the unique alignment of fibres. A quantitative understanding of the topological order that governs the orientational field of cardiac fibres is essential for elucidating the mechanical basis of cardiac performance.

In recent years, concepts from liquid crystal physics have emerged as a tool to describe orientational order in biological tissues [8]. Nematic order (Fig. 1a), characterised by a long-range orientational order without positional order of its constituent particles, has been shown to play critical roles in cell culture, epithelial monolayers, and tissue morphogenesis [9–11]. Two-dimensional (2D) studies have revealed that topological defects in nematic fields are associated with self-propulsion [12], function as sources and sinks of cell flow [13] as well as cell death [14], or precede organ regeneration [15]. However, biological tissues are inherently 3D, and the full richness of 3D nematic topology remains largely unexplored due to experimental and analytical challenges, apart from a few notable exceptions that have analysed purified microtubules [16] and malignant tissues [17].

**Fig. 1.**
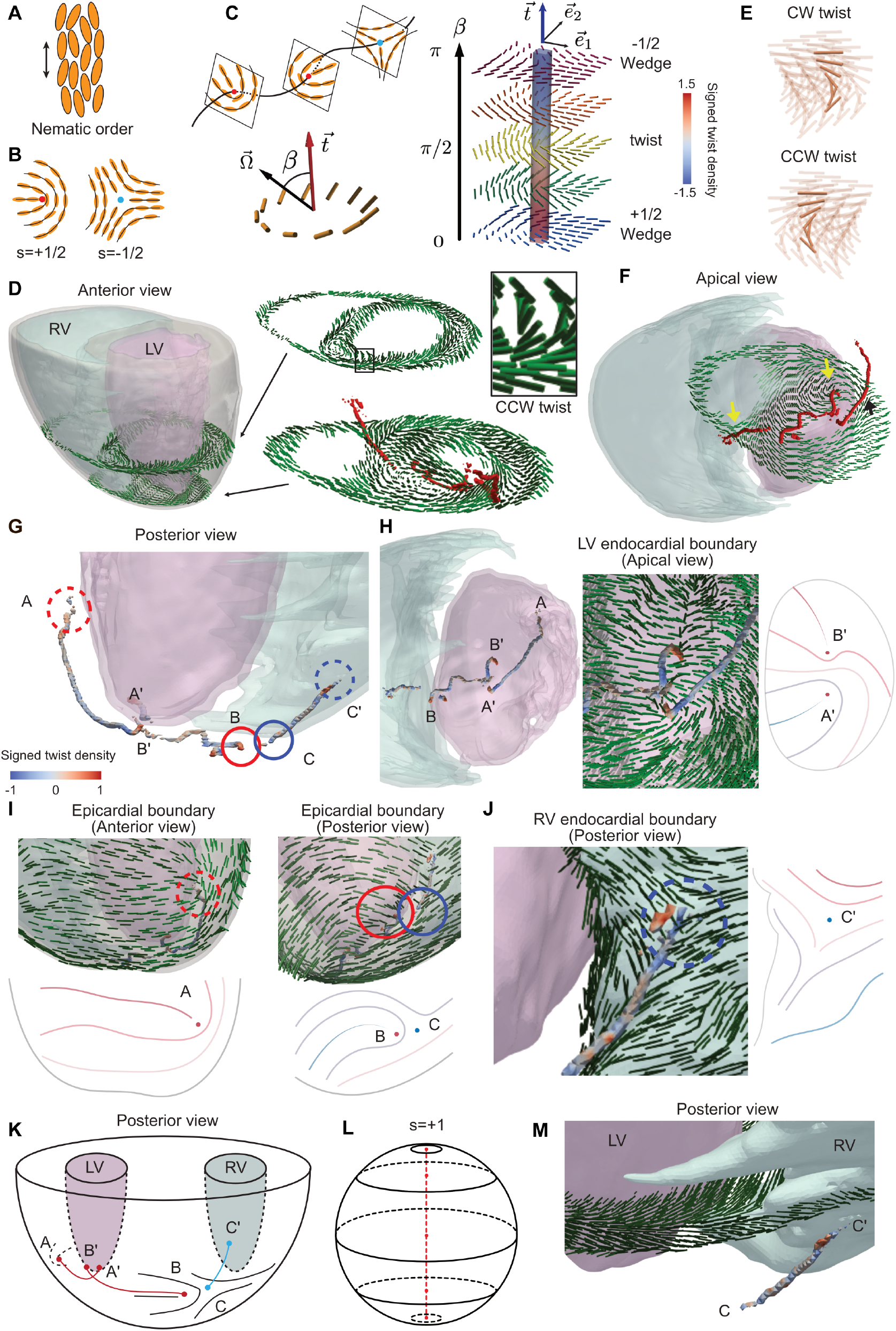
Topological characterisation of 3D fibre architecture in the adult mouse heart. (a) Orientational order in nematic materials. (b) Topological defects in two dimensions, characterised by half-integer topological charges. (c) In 3D, topological defects appear as disclination lines. The local pattern surrounding a disclination line is characterised by the angle β between the tangent direction of the line 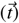 and the rotation axis 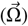 around the line. The colour of the disclination line represents the signed twist density (the trace of the disclination density tensor). (d) Three-dimensional nematic pattern of cardiac fibre structure in the adult mouse heart, showing CCW transmural twist and disclination lines. (e) Three-dimensional spatial twist exhibits distinct chirality, classified as CW or CCW. (f) Apical view of a cross-section of the orientation field with disclination lines. Two-dimensional defect-like regions are indicated by arrows. (g) Magnified view of disclination lines from the posterior side. Heat colour represents the signed twist density. (h) Orientation field surrounding disclination line endpoints in LV. (i) Sub-epicardial orientation pattern in the vicinity of disclination line endpoints. (j) Magnified view of the endocardial endpoint of the disclination line on RV. (k) Schematic representation of disclination lines within the cardiac fibre architecture. (I) Schematic of a typical tangential orientation field and topological defects on a spherical surface. (m) Magnified view of the orientation field close to the sharp edge of RV endocardial boundary. CCW, counter-clockwise; CW, clockwise; LV, left ventricle; RV, right ventricle.

The myocardium is a paradigmatic example where 3D nematic order and biological function intersect: cardiac fibres form a chiral nematic field [8] and spontaneously contract to achieve coordinated contraction. While topological defects have been identified in 2D sections of the heart [18, 19], their existence, geometry, and functional significance in the 3D cardiac tissues are unknown. Addressing this gap is both a fundamental challenge and an opportunity for new biomechanical principles relevant to health and disease.

In this work, we treat myocardial fibre orientation as a coherent chiral nematic field, independent of discrete fibre bundles, enabling a topological description of cardiac architecture in toto. Using diffusion tensor MRI data and high-resolution light-sheet imaging, we developed a method to quantify the orientation pattern and identify disclination lines (i.e., topological defects in 3D nematics). This method was applied to the normal mouse heart and the heterotaxy (left-right inverted) mouse heart to show that tissue chirality is independent of systemic left-right signaling or global heart geometry. Mechanical simulations further indicate that these nematic architectures support contractile function, with local tissues near topological defects exhibiting reduced work output and global contractile performance depending on coherence of myocardial fibre chirality. By viewing the heart as a chiral nematic field, our results uncover how local twists are integrated into a global structural organisation, and provide insights into the biomechanical principles underlying cardiac performance and morphogenesis in the complex multicellular system.

## Identification of 3D topological defects in adult mouse heart

Nematic order describes the collective alignment of rodshaped entities such as in liquid crystals (Fig. 1a). The director field can bend or splay in 2D, and topological defects appear when the orientation is discontinuous. These point defects are characterised by a half-integer winding number *s*. The director rotates through an angle 2π*s* when one traverses a closed loop around the defect [8]. The simplest examples are the wedge defects with *s* = ±1/2 (Fig. 1b).

In 3D, defects can take the form of lines, called disclination lines (Fig. 1c). Compared with the 2D case, which can exhibit infinitely varied types of half-integer defects, the 3D case has only a single type of line defect [20, 21]. Cutting a disclination with any plane shows that the director rotates by π around the intersection point; however, the axis of this rotation can vary continuously along the line. Consequently, a wedge +1/2 profile seen in one cross-section can smoothly transform into a wedge −1/2 profile in another. This spatial variation can be quantified by the local rotation axis Ω and the twist angle β, defined as the angle between Ω and the tangent vector ***t*** of the line. By convention, a wedge +1/2 defect has β = 0, a wedge −1/2 defect has β = π, and a configuration with β = π/2 is called a twist profile (Fig. 1c).

Disclination lines can be identified as the topological invariant of the disclination density tensor ***D*** in 3D [22]. Regions with a large Frobenius norm of ***D*** correspond to the core of the disclination line. The trace of ***D***, referred to as the “signed twist density”, is proportional to cos β and large positive (negative) value indicates a wedge +1/2(−1/2) defect type (see Methods). The second-rank disclination density tensor ***D*** can be expressed in terms of the nematic order parameter ***Q***:

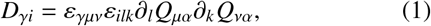

where ε_*i jk*_ is a Levi-Civita symbol and ∂_*i*_ represents a spatial partial derivative [22]. The subscripts denote 3D Cartesian components of the tensors.

Using Equation (1), we established a computational framework to analyse the orientation order and disclination lines in the mouse heart from diffusion tensor imaging (DTI) data obtained by Angeli et al. [23] (see Methods). The disclination density tensor ***D*** was computed from the fibre orientation field reconstructed from the DTI data. Then, disclination lines were extracted as one-dimensional ridges of high Frobenius norm values of ***D*** (Fig. 1d and Extended Data Fig. 1). DTI images of adult mouse hearts revealed a chiral orientation field exhibiting a CCW transmural twist, together with disclination lines displaying distinct spatial patterns within the ventricular wall (Fig. 1d,e). The local nematic field profiles resembled the 2D topological defects in the short-axis cross-sectional planes (Fig. 1b,f) [24].

Extensive analysis revealed three major core disclination lines A-A’, B-B’ and C-C’ in the murine heart (Fig. 1g). Disclination line A-A’ extended from the apex of the epicardial boundary (A) to the endocardial boundary (A’), with both endpoints carrying +1/2 surface charge (Fig. 1h and 1i). Disclination line B-B’ connected the epicardial apex and the endocardial boundary of the RV (Fig. 1g-i), and its points also exhibited +1/2 wedge surface defects (Fig. 1h). Despite the complex fine structure of the posterior endpoint (B), the net surface charge remained +1/2 (Fig. 1i). The third disclination line C-C’ originated from the posterior epicardial boundary and resolves at the RV endocardial boundary, with both endpoints exhibiting −1/2 wedge surface defects (Fig. 1j). The whole configuration of the disclination lines is depicted in Fig. 1k. Comparable patterns were observed in two additional samples, supporting the presence of a common topological structure, while revealing variability in whether the major disclination lines, B-B’ and C-C’, are connected or not (Extended Data Figs. 2 and 3).

The topology of the disclination lines is constrained by the director orientation on the ventricular and epicardial surfaces. For example, on a spherical surface with tangential director orientation, the total topological charge is fixed to +2, typically appearing as two +1 surface defects on the north and south poles (Fig. 1l). Consistent with this constraint, the LV endocardial boundary in the adult murine heart exhibited nearly tangential alignment and contained two +1/2 surface defects, totalling +1 charge at the apical pole (points A’ and B’ in Fig. 1k, Extended Data Figs. 2 and 3). In contrast, the anterior and posterior RV-septum insertions form sharp edges, which allows the surface orientation to be perpendicular at the edges (Fig. 1m). Due to this broken tangential orientation and the thin RV free wall, the structure of the surface defects on the RV endocardial and epicardial boundaries could not be reliably resolved.

The fibre topology was also analysed in embryonic hearts at E18.5 using confocal microscopy images (Extended Data Fig. 4 and Methods). Similar to the adult heart, a disclination line was identified at the apex (A-A’). The disclination lines B-B’ and C-C’ in adult samples (Extended Data Figs. 2 and 3) were often connected into a single line (B’-C’) in the embryos. Besides these similarities, there were also significant differences in the distribution of disclination lines between embryonic and adult hearts. First, the endpoint C’ resolved at the basal epicardial boundary instead of the endocardial boundary of the RV. Second, two additional disclination lines extended from the endocardial boundary of the RV to the epicardial boundary (D-D’ and E-E’). Therefore, the topological structure in the cardiac fibre may undergo remodelling during perinatal and postnatal growth.

Together, these observations suggest that the cardiac fibres exhibit chiral nematic order with disclination lines extending from the apex with variability across individual animals and maturational stages.

## Distinct mechanical properties of cardiac tissue near topological defects

The active nematic theory has demonstrated that regions close to topological defects in 2D cell monolayers may experience anomalous stress and exhibit distinct mechanical properties [12–15]. Therefore, to analyse sarcomere dynamics in a beating heart with anisotropic fibre structure, we next examined the mechanical properties of topological defects using an established mechanical simulation pipeline that incorporates molecular-level state transitions [25–27]. A finite element model based on the actual organ geometry and fibre orientation was constructed from the same mouse DTI data as in Fig. 1 (Fig. 2a and Extended Data Fig. 5, see Methods.)

**Fig. 2.**
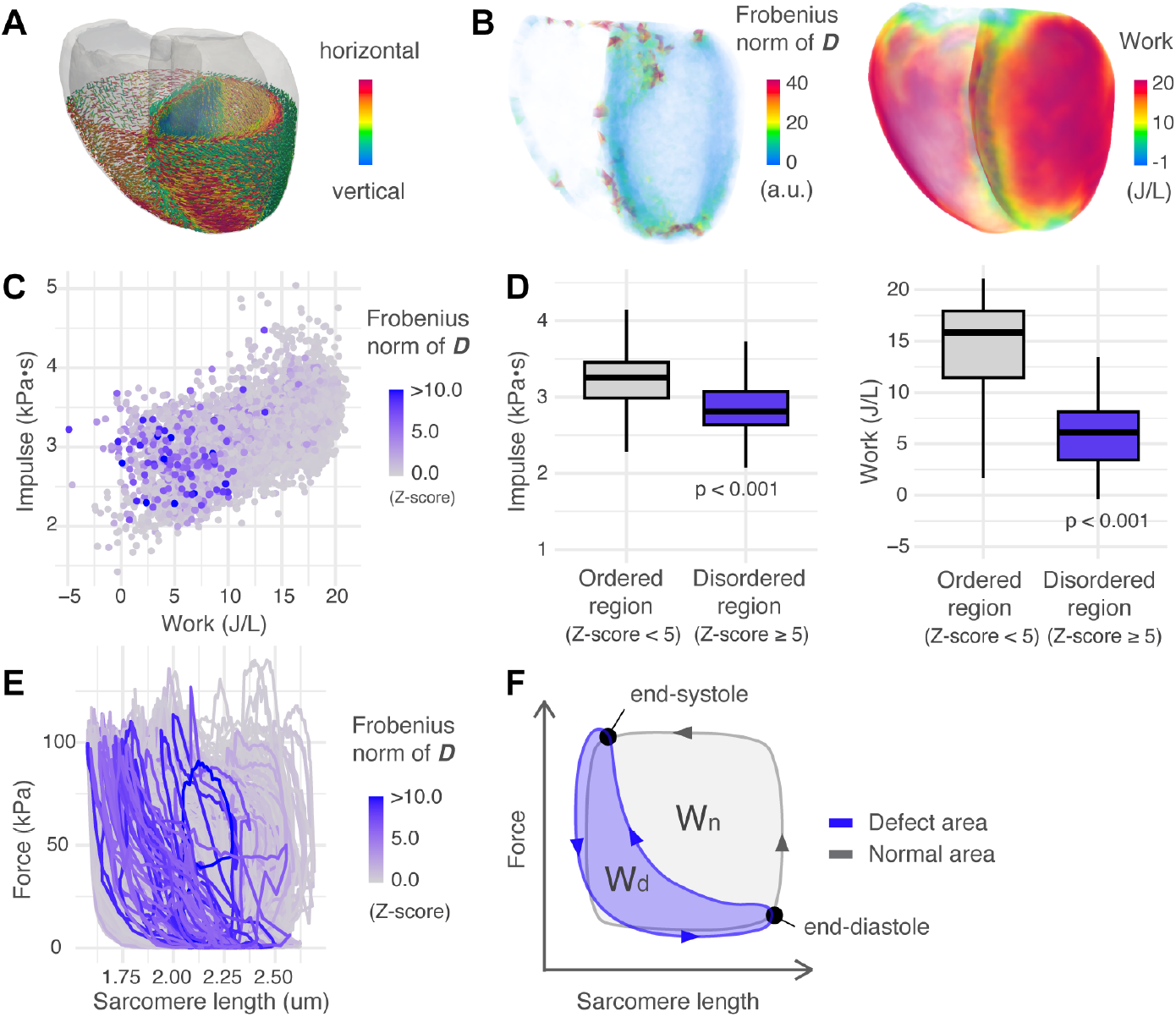
Topological defects exhibit impaired local mechanical efficiency in three-dimensional cardiac tissue. (a) Finite element method (FEM) mechanical simulation model. The simulation models assume fibre orientation based on diffusion tensor imaging data and implement anisotropic contraction along the local fibre direction. (b) Spatial distribution of topological defects and work generation in cardiac tissue. Defect regions were identified as areas exhibiting high values of the disclination density tensor norm (left). Work generated over one cardiac cycle was reduced in regions coinciding with disclination lines (right). (c) Impulse and work generation in individual FEM elements. Colour represents the z-score of the Frobenius norm for each element. High z-scores indicate large values of the Frobenius norm caused by irregular fibre alignment. Each dot represents a single FEM element. (d) Box plots of impulse and work in tissue with well-aligned or irregular fibre structure. Regions corresponding to disclination lines were distinguished by high z-scores (≥5) and compared with other regions. Both impulse and work were significantly reduced in high z-score regions. P-values were calculated using Welch’s two-sample *t*-test. (e,f) Sarcomere length-force relationship during the cardiac cycle and its schematic representation. Due to the early completion of contraction, the area of the loop, corresponding to mechanical work, was reduced in defect regions.

We observed that both the work and impulse generated by the tissue were significantly reduced in the vicinity of topological defects (Fig. 2b-d). To elucidate the underlying mechanics, we resolved force generation and sarcomere length throughout the cardiac cycle. While local force generation remained comparable to that of defect-free tissue, sarcomeres exhibited rapid shortening during the contraction phase (Fig. 2e). This suggests that the discontinuity in fibre orientation effectively uncouples local contractile forces from the mechanical traction of the surrounding tissue. Consequently, the tissue undergoes rapid contraction against reduced resistance, leading to the observed diminution in work output (Fig. 2f).

## Tissue-intrinsic chirality of fibre structure at the apex

In heterotaxy syndromes, the left-right (L/R) body plan is inverted, leading to the mirror image positioning of visceral organs, including the heart. However, earlier anatomical studies have reported that the orientation of sub-epicardial muscle fibres remains normal in patients with heterotaxy [6, 28, 29], pointing toward the possibility that myocardial fibre chirality may be independent of systemic L/R cues. To test this, the myocardial fibre architecture of *inv*/*inv* mice at stage E18.5 was compared with that of control littermates, a murine model of situs inversus totalis (SIT) caused by loss-of-function mutations in the inversin (*inv*) gene [30, 31].

The macroscopic cardiac structures (organ morphology, atria, ventricles, large vessels, and coronary arteries) exhibited mirror-symmetric organisation in inverted hearts compared with normal hearts (Fig. 3a, Extended Data Fig. 6). The arrangement of disclination lines also showed structures corresponding to nearly mirror-image positions (Fig. 3b).

**Fig. 3.**
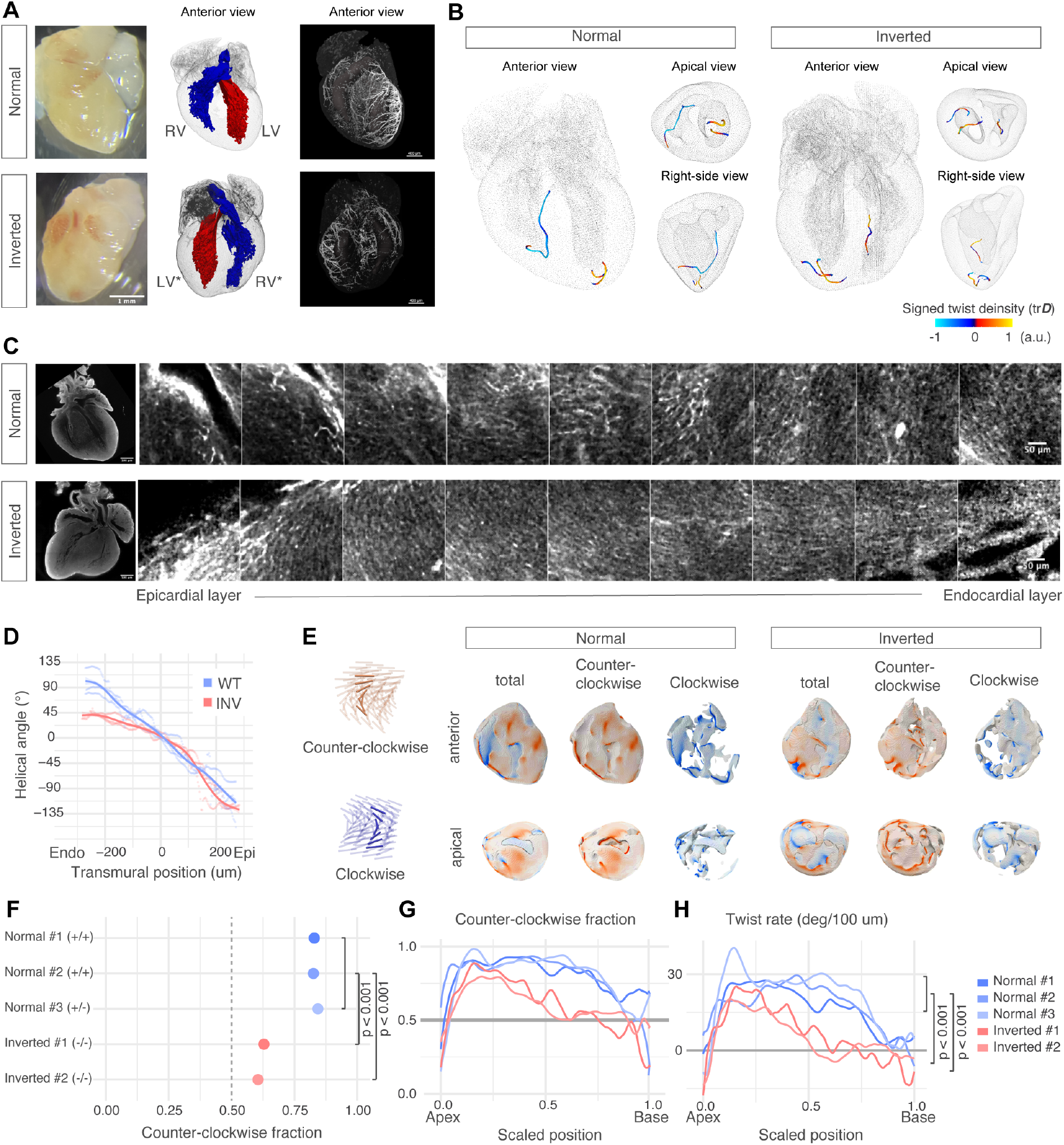
Preserved CCW chirality in heterotaxy hearts despite global left-right inversion. (a) Mirrored macroscopic structures of a heterotaxy heart at E18.5. The inverted heart exhibits mirror-symmetric organisation of overall morphology, including ventricular and atrial outflow tracts and coronary arteries, relative to the normal heart. (b) Disclination lines are detected at mirror-symmetric positions in normal and inverted hearts. Colour indicates the trace of the disclination density tensor D. (c) Confocal microscopy images from epicardial to endocardial layers showing helical structure in the anterior wall of the left ventricle in normal and inverted hearts. (d) Quantification of helical angle across the ventricular wall. (e) Three-dimensional tissue chirality of cardiac fibre structure. Colour represents the magnitude of twist density. Both normal and inverted hearts demonstrate predominantly CCW chirality throughout the majority of the tissue. (f) Volume fraction of tissue exhibiting CCW chirality in each organ. Despite whole-body left-right signal inversion and macroscopic morphological mirroring, inverted hearts retain CCW chirality. P-values were calculated using a one-sided t-test (n = 5). (g, h) Profiles of CCW chirality and twist rate along the apex-base axis. Normal hearts exhibit consistent CCW chirality throughout the entire organ. Inverted hearts display clear CCW chirality in the apical region, while the basal region contains intermixed CW domains. P-values were calculated using a one-sided t-test (n = 5). CCW, counter-clockwise; CW, clockwise; LV, left ventricle; RV, right ventricle; LV*, anatomical left ventricle; RV*, anatomical right ventricle.

To investigate the tissue chirality of the inverted heart, we first measured the 2D helical angle. A CCW twist was observed from the epicardium to the endocardium in the inverted hearts, similarly to the normal hearts. In inverted hearts, however, the twist was weaker towards the endocardium (Fig. 3c,d). These findings are consistent with previous reports [32].

To examine the chirality of the inverted heart in 3D, we quantified the twist density *q* = ε_*i jk*_*Q*_*il*_∂ _*j*_*Q*_*kl*_ (see Methods). The volume fraction of CCW-oriented myocardium was around 0.8 in normal hearts (Fig. 3e). However, inverted hearts exhibited markedly reduced CCW-oriented component (volume fraction closer to 0.5, Fig. 3f and Extended Data Fig. 7a). Moreover, the position-wise volume ratio and mean twist rates were calculated to examine chirality in greater detail. The results revealed that normal hearts consistently exhibit CCW chirality continuously from the apex to the base. Conversely, the SIT hearts retained CCW dominance in regions near the apex while progressively gaining a CW component towards the base (Fig. 3g, h and Extended Data Fig. 7b, c). Thus, although the global geometry of the inverted heart is a mirror image of the normal heart, the twist direction is chimeric, with an uninverted (CCW) twist in the apical side of the heart.

The chiral structure of myocardial fibres is considered to be influenced by cell intrinsic chirality, external signalling cues, and constraints imposed by boundary conditions. The observation that even inverted hearts retain CCW chirality suggests a minimal direct contribution from systemic L/R signal to the twisting in the tissue. Moreover, the well-preserved CCW chirality at the apex, a region less constrained by boundary conditions from surrounding tissues, suggests that myocardial cells possess intrinsic chirality capable of generating CCW twisting. In contrast, the reduced fibre twist observed at the base region of inverted hearts may reflect the stronger influence of reversed boundary constraints imposed by surrounding tissues.

## Consistent fibre chirality is crucial in cardiac function

The chimeric chirality in inverted hearts provides a valuable model for studying the role of transmural fibre twist in the cardiac pump function. To explore this, we first examined how the cardiac fibre chirality affects cardiac function. Artificial orientation fields were generated using an impulse-maximization optimization method as we previously reported [25–27]. This process amplifies initially minimal helical orientations into well-developed transmural twists, enabling us to create structures with CCW twist (control), no twist, and complete CW twist (artificial) for simulation within the standard heart shape (Fig. 4a and Extended Data Fig. 8 and 9). The beating simulation demonstrated that fibre structures with twists, both CCW and complete CW, exhibited better cardiac function than those without twists. Structures without twist exhibited reduced efficiency in ejection fraction, maximum pressure, and stroke volume, revealing impaired ejection performance due to the lack of omnidirectional contraction (Fig. 4b).

**Fig. 4.**
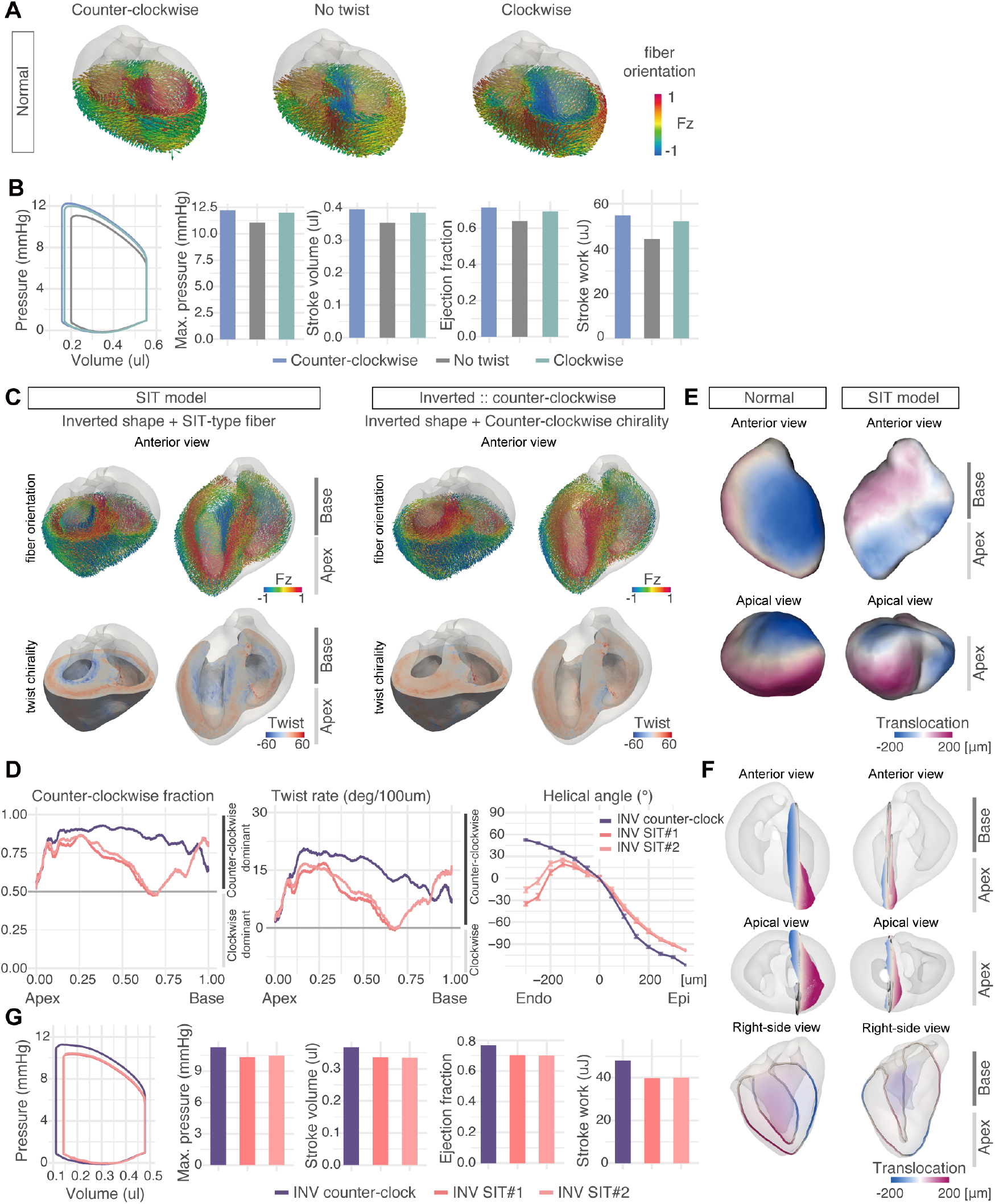
Tissue chirality and organ function. (a) CCW or CW twist structure with a normal heart shape. Fz indicates the element of the unit vector in the apical-to-basal direction. (b) Pressure-volume curves and heart functions with different twist chirality. (c) Fibre structures of the situs inversus model (SIT) and control with an inverted heart shape. The SIT model has a mixture of CCW twist in the apex and CW twist in the basal region, whereas the control heart has a consistent CCW twist structure. Colour indicates fibre orientation or twist rate (degrees per 100 µm). (d) Profiles of chirality and the twist rate of the inverted heart simulation model in the apex-to-base direction (left) or endocardial-to-epicardial direction (right). (e, f) Cardiac motions of the SIT model. Overall (e) and cross-sectional (f) views. Colour indicates the left-right direction and the distance of local tissues in the contraction phase from their end-diastolic position. (g) Pressure-volume curves and heart functions with CCW or mixed chiral structure.

Intriguingly, mechanical simulation revealed no differences between the CCW and artificially-generated complete CW twist models. While hearts without twist can contract circumferentially, impaired longitudinal contraction leads to abnormal protrusion at the apex during systole (Fig. 4b) [27]. Thus, the overall pumping function depends not on the chirality itself but on the consistency of chirality throughout the myocardium.

To further investigate how heterogeneous twist affects the cardiac contractile motion, we generated models based on the SIT fibre structure, characterised by a CCW twist at the apex and CW twist in the interior near the base [33]. A structure with a uniformly CCW twist was also prepared using the same inverted heart geometry (Fig. 4c). These structures exhibited chirality and twist rate profiles similar to actual measurements (Fig. 4d).

Lastly, we conducted finite element simulations to assess the effect of chimeric chirality on the contractile performance. Hearts with the SIT configuration exhibited torsional motion congruent with controls at the apex but opposing at the base (Fig. 4e, f, Extended Data Fig. 10 and Extended Data Movie 1), consistent with previous observations [33, 34]. Quantitatively, the SIT heart displayed reductions in maximum pressure, stroke volume, ejection, and stroke work relative to the uniformly twisted model (Fig. 4g). These findings demonstrate that a heterogeneous twist compromises cardiac performance, providing direct evidence that the homogeneously twisted topology of cardiac fibre is crucial for cardiac contractile function.

## Discussion and conclusion

Our analysis of heterotaxy hearts uncovers that two distinct but interacting chirality underlie cardiac morphogenesis. Global left-right signalling dictates large-scale asymmetry, such as the arrangement of ventricles and atria as well as the overall heart shape, whereas the local helical structures of cardiomyocytes generate consistent chiral fibre organisation, particularly at the apex and sub-epicardial layers. The coexistence of these independent mechanisms and the partial inversion observed at the base of heterotaxy hearts illustrates how global patterning cues and local self-organisation jointly form the heart’s structure. This interplay may represent a general principle of organogenesis, in which intrinsic tissue chirality interacts with global developmental axes.

This dissociation is exemplified by the *Inv*/*Inv* mouse model. Although this strain carries a mutation in a cilia-associated protein that reverses whole-body left-right asymmetry, the myocardial fibre architecture retains its characteristic CCW chirality. This confirms that local myocardial organisation is robust to the reversal of global nodal flow. Consequently, our mechanical simulations demonstrate that the modest functional deficits observed in *Inv*/*Inv* hearts can be explained solely by the resulting mismatch in local nematic structure (chimeric chirality), without invoking additional cilia-derived cellular defects.

The molecular origin of this cell-intrinsic chirality is potentially the actomyosin cytoskeleton. A broad survey of micropatterned mammalian cell types revealed phenotype-specific left–right rotational biases that depend primarily on actin rather than microtubule integrity [35]. In fibroblasts, this chirality arises through myosin-IIA-driven tilting of formin-nucleated radial actin fibres [36], while in epithelial cells it emerges from myosin II torque acting on a dorsal concentric actomyosin network, driving directional nuclear rotation without requiring cell-scale chiral cytoskeletal order [37]. In *C. elegans*, active torques generated by the actomyosin cortex contribute to symmetry breaking at the embryonic scale [38]. Cardiomyocytes themselves exhibit cell-autonomous chiral behaviour: isolated chick cardiomyocytes rotate with a preferred handedness in vitro, driven by N-cadherin and asymmetric phospho-myosin II [39, 40], providing a plausible actomyosin-based cellular origin for the CCW transmural fibre twist.

Crucially, this fibre twist chirality is mechanistically distinct from organ-level cardiac looping: while looping is reversed in inv/inv hearts under global left-right inversion, the transmural fibre twist is preserved, indicating it is encoded at the cytoskeletal level and largely insulated from systemic left-right patterning signals. This dissociation carries a fundamental implication: for normal cardiac function, the specification of left-right directionality at the organ level must be compatible with the cell-intrinsic chirality of cardiomyocytes, a constraint that, when violated, produces the chimeric fibre organisation and mechanical impairment seen in heterotaxy.

The nematic order of myocardial fibres provides mechanical advantages that enhance cardiac efficiency. As we observed through simulations, regions with coherent fibre alignment generate greater contractile work, while local distortion reduces mechanical output. The transmural twist of fibre orientation enables coordinated circumferential and longitudinal contraction, supporting effective blood ejection. Hearts with inconsistent twist patterns exhibit impaired performance, indicating that consistency of the twist chirality, rather than the chirality per se, is critical for efficient pumping function. This mirrors what is clinically seen at the level of organ placement: complete left–right inversion is often well-tolerated, whereas mixed chirality leads to substantial dysfunction.

From the perspective of nematic topology, the long-debated helical ventricular myocardial band [41–43] can be interpreted as a consequence of the ordered fibre field, enabling the tracing of a continuous band-like pathway from the pulmonary artery to the aorta without encountering topological defects. However, this band is achieved through selective dissection of the left ventricle and septum into discrete layers, even though fibre orientations exhibit continuous CCW transmural twist across these layers [4]. This suggests that while the helical band concept captures certain organizational principles, the nematic-order description may offer a more general and continuous characterization of myocardial fibre architecture.

Understanding the topological organisation of cardiac fibres has implications for both basic biology and clinical medicine. Disorders in fibre alignment, as seen in hypertrophic and dilated cardiomyopathy, may involve disruptions to the nematic order, while topological defects could create electrophysiological substrates that promote re-entrant circuits in ventricular arrhythmias. Furthermore, alterations in transmural twist patterns may underlie the mechanical dysfunction characteristic of heart failure with preserved ejection fraction. Recent advances in cardiac organoid technology and computational modelling have highlighted the importance of fibre orientation in engineered cardiac tissues [44, 45], yet the fundamental topological principles governing this organisation have remained elusive. Our findings provide a theoretical foundation that could inform novel strategies in both tissue engineering approaches for regenerative therapies and in silico modelling for drug discovery and personalised medicine.

## Methods

### Animal treatments

All animal procedures in this study were approved by the UCLA Animal Research Committee (ARC)-2008-126, and this study complied with all relevant ethical regulations while conducting animal experiments.

### MRI data acquisition

Adult mouse MRI data were obtained from a previously published dataset by Angeli et al. [23]. Diffusion tensor imaging data from three 8-12-week-old male C57BL/6 mice were analyzed using the acquisition parameters described in the original study.

### Mouse sample preparation

Healthy mice (*inv*/+) were set up for timed mating, and embryos were collected at embryonic day 18.5 (E18.5). Heart samples were collected and fixed with 4% paraformaldehyde/phosphate-buffered saline (PBS) overnight. Littermate pairs consisting of one normal (WT or *inv*/+) and one inverted *(inv*/*inv)* mouse from three different breeding pairs were examined. Inclusion criteria required intact ventricular morphology without gross developmental abnormalities, while exclusion criteria eliminated samples with imaging artifacts or incomplete tissue clearing.

### Whole mount staining and tissue clearing of embryonic mouse heart

Fixed heart samples were washed with PBS extensively 1 day before further treatment. They were treated with 5% H_2_O_2_/MeOH overnight at 4°C to reduce auto fluorescence in cardiomyocytes and blood cells. CUBIC-L/R (TCI, T3740 and T3983) or RapiClear (SUN Jin lab, RC152001) was used for tissue clearing; samples were incubated with CUBIC-L or RapiClear at 37°C for 0.5–7 days until they became transparent. Cell membrane was visualised with WGA Alexa-647 (Thermo Fisher, W32466), and SYTOX Green (Thermo Fisher, S7020, 1:100 dilution in 500 mM NaCl/PBS) or RedDot2 (Biotium, 40061, 1:100 dilution in 500 mM NaCl/PBS) was used for counterstaining of nuclei. Staining was performed at room temperature for 0.5– 2 day(s) according to the size of the sample. After extensive washing with PBS, samples were incubated in CUBIC-R solution at room temperature for RI matching.

### Image acquisition with confocal and light-sheet microscopy

Confocal imaging was performed using a Zeiss LSM880 inverted microscope equipped with an objective lens (Plan-Apochromat 10×/0.45) and lasers (488, 561 and 633nm). Cleared heart samples were embedded in 10% Agarose (Nacalai Tesque, 01163-76) in CUBIC-R solution. The samples were imaged with Z stacks of optical sectional images with 0.6× –1.0× digital zoom (200–300 slices with an interval of 3.05 um). For lightsheet imaging, a Zeiss LS7 microscope with a 5× objective lens and lasers (488, 561 and 647 nm) was used. Heart samples were embedded in 10% Agarose/CUBIC-R in a glass cylinder 1 day before imaging, and placed in the mounting chamber filled with the mounting solution for CUBIC-R (TCI, M3294). 3D image stacks were obtained (1440×1440 pixels, 2.88 um step interval). XY tiling was used for large samples to cover the entire heart region, and tiles were stitched using Fiji/ImageJ [46–48] and Imaris stitcher (Bitplane, version 10.0.0).

### Image analysis of microscopy and MRI data

The 3D orientation field was derived from microscopy images and MRI data. For microscopy images, the structure tensor ***J*** was calculated using python structure-tensor package [49] based on the following equation:

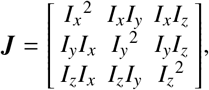

where *I*(*x, y, z*) is the intensity at each voxel, *I*_*x*_ = ⟨∂_*x*_*I*(*x, y, z*)⟩ with ∂_*x*_ denoting the partial derivative in the *x* direction, and the brackets ⟨·⟩ indicating local spatial averaging. The parameters for the spatial averaging implemented in the structure tensor 3d function were set as σ = 1.0, ρ = 10.0. The principal axis ***n*** of ***J*** was first computed, and the nematic tensor was reconstructed via *Q*_*i j*_ = 3/2(*n*_*i*_*n*_*j*_ −δ_*i j*_/3).

For MRI data, the 3D nematic tensor *Q*_*i j*_ was calculated from the diffusion tensor ***d***. To reduce the noise, the principal axis ***n*** of ***d*** was first computed, and the nematic tensor was reconstructed via 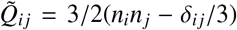. The component of 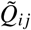 was then denoised using a (5 pixels)^3^ uniform mean filter. The principal axis ***ñ*** was extracted again from 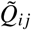, and a filtered nematic tensor 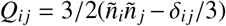 was reconstructed.

### Detection of 3D disclination line

The disclination lines were detected from the orientation field following the method proposed by Schimming et al. [22]. In brief, the disclination density tensor ***D*** is defined by Eq. 1. Since this tensor is constructed from the spatial variations of the order parameter, when the orientation field is aligned parallel, ***D*** equals zero. In contrast, the elements of ***D*** take on large values around singular points. Thus, the topological defect was detected as the region where ***D*** has a significantly large Frobenius norm, defined as follows:

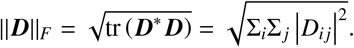

This approach allows for the precise identi fica tion of regions where the orientation field exhibits significant discontinuities or singularities, which effectively maps the disclination lines in the volume images of the heart.

### 3D tissue chirality

The chiral order parameter, or twist density, measures the 3D tissue chirality:

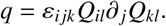

whose positive or negative sign corresponds to CCW or CW rotation. A more intuitive description of *q* is provided as *q* = (9/4)(***n*** ·∇×***n***) = − (9/4)∂_*z*_θ, where ***n*** is a vector representation of the fibre orientation, and the local frame is provided as ***n*** = (cos θ, sin θ, 0). Using a right-handed coordinate system, *q* > 0 corresponds to the CCW chirality. Thus, the twist rate was calculated by 4*q*/9, and the CCW volume fraction was defined as the ratio of volume where *q* > 0 to the total volume.

### Mechanical modelling and simulation

Mechanical simulation for functional analysis of the heart with fibre orientation field was performed based on previously published multiscale electromechanical heart models [25–27]. This framework has been shown to quantitatively reproduce key features of cardiac pumping in the human heart, including realistic ventricular pressure-volume dynamics, spatially heterogeneous strain patterns, and physiologically consistent timing of contraction and relaxation, by explicitly coupling molecular-scale force generation to tissue-scale mechanics.

Briefly, cardiac contraction was driven by a prescribed time-dependent intracellular calcium concentration waveform, which regulates the stochastic attachment-detachment dynamics of myosin cross-bridges in a half-sarcomere model embedded within each finite element. For a given calcium transient, the cross-bridge model computes the active tension generated along the local myofibre direction, incorporating force-length and force-velocity effects arising from filament sliding. The resulting active tension was incorporated into the continuum description of the myocardium as an anisotropic active stress aligned with the fibre direction and combined with passive hyperelastic and viscous stresses. The equations of motion for the myocardial tissue were solved using a finite-element formulation.

Local fibre stretch and shortening velocity, determined from the deformation field, were fed back to the cross-bridge model, thereby establishing a bidirectional coupling between molecular-scale force generation and tissue-scale mechanics. Mechanical interactions between neighbouring finite elements were mediated through the global displacement field, allowing local contraction to influence loading and deformation in surrounding regions. Within this framework, the time courses of fibre stretch (and corresponding sarcomere length) and active tension were obtained for each finite element, enabling direct construction of sarcomere length-force relationships. Mechanical work was calculated by integrating the product of active tension and fibre shortening over time, either locally at the element level or globally over the ventricular wall.

Cardiac geometry was extracted from microscopy images or MRI data using Imaris software (Bitplane, version 10.0.0) to isolate the shape of the myocardium. Gmsh software [50, 51] (version 4.12.2) was then used to create the tetrahedral mesh model necessary for the finite-element simulations. The fibre orientation field was assigned based on diffusion tensor values evaluated at the centre of gravity of each element, derived from diffusion tensor MRI data.

Cardiac beating simulations were performed by modifying the human beating conditions from the previous study to match mouse conditions. For the adult mouse, the heart rate was set to 440 beats per minute, and all state transition rates applied in the human heart model (HR= 60) were multiplied by a factor of 440/60 = 7.3. For the embryonic mouse, the heart rate was set to 140 beats per minute, and all state transition rates were multiplied by 140/60 = 1.3. Additionally, it was necessary to reduce contractile force to reproduce the blood pressure values to be lower than normal. The proportion of myofibrils with contractile capacity was set to 50% of normal, and intracellular calcium ion concentration was also set to 0.8 times the normal level.

### Generation of cardiac fibre orientation field with a distinct chirality

Cardiac fibre structures with various chiralities in the embryonic heart were generated following previous studies [25, 27]. The initial orientation before applying optimisation was determined as a function of location using the helix angle α according to the following equation:

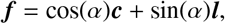

where ***c*** is the unit vector in the circumferential direction and ***l*** is the unit vector in the longitudinal direction. The initial orientation was determined by defining the helix angles α_*endo*_ and α_*epi*_ at the endocardium and epicardium, respectively, and linearly interpolating in the transmural direction as follows: CCW: α_*endo*_ = 10°, α_*epi*_ = −10°, No twist: α_*endo*_ = 0°, α_*epi*_ = 0°, CW: α_*endo*_ = −10°, α_*epi*_ = 10°. For the SIT model, α values were determined for each height *h* from apex to base at the endocardium and epicardium, and for *h* ≥ *h*_*M*_, an additional midpoint α was defined by linear interpolation with respect to *h*, followed by transmural interpolation. SIT#1 and SIT#2 differed only in *h*_*M*_; SIT#1: *h*_*M*_ = 0.25, SIT#2: *h*_*M*_ = 0.4 (see Extended Data Fig. 8).

The optimisation of the orientation field ***f*** was then executed according to the maximising impulse with modified parameters for the mouse heart (weak contractile force 4 kPa, cone angle 30°), and the process was terminated after 200 steps when the intraventricular pressure reached a plateau.

### Statistics

Statistical significance was examined by custom R coding (R 4.4.3). Welch’s two-sample *t*-test (ggpubr::stat compare means, tidyverse 2.0.0) was used in Fig. 2, or one-sided *t*-test (t.test in base package) in Fig. 3. The null hypothesis in Fig. 3 is that the observed value from the inverted heart is the same as the mean from standard heart samples. The profile and the apex-base axis were examined at 0.6 in the scaled position.

## Supplementary information

## Acknowledgements

We are deeply grateful to Dr. Stelios Angeli and Dr. Christakis Constantinides for generously providing the mouse MRI data, originally acquired and provided by the Center for In Vivo Microscopy, Duke University. We sincerely appreciate their willingness to allow us to use their individual data sets in this research. The confocal microscopy was performed using the facilities at the UCLA Microscopy Core.

This work was supported by the following grants; N.K. is funded by Takeda Science Foundation Oversea fellowship, T.Y. is funded by RIKEN Stage Transition Project.

## Author contributions

N.K. and T.Y. conceptualised the examination of nematic order in cardiac fibres. N.K. and A.N. conceptualised examining the functional significance of the fibre topology. N.K., T.Y., K.K. and A.N. designed the study. N.K. performed all the experimental procedures using mouse heart tissue and analysed the microscopy data. T.Y. and K.K. developed the theoretical methodology and analysed the MRI data. T.W. and N.K. performed the mechanical simulation and analyses. V.X. contributed to light-sheet microscopy observations. Y.N. and K.Y. set up *inv*/*inv* mice breeding and collected heart samples. All the authors contributed to the discussion of the results. N.K., T.Y., K.K. and A.N. wrote and edited the original manuscript.

## Data Availability

The datasets generated and analysed during the current study are available from the corresponding author upon reasonable request. The adult murine MRI data used in this study are those used in [23]; additional permission must be obtained directly from the authors for data access and reuse.

## Code Availability

The custom code used in this study is available from the corresponding author upon reasonable request.

---

**Extended Data Fig. 1.**
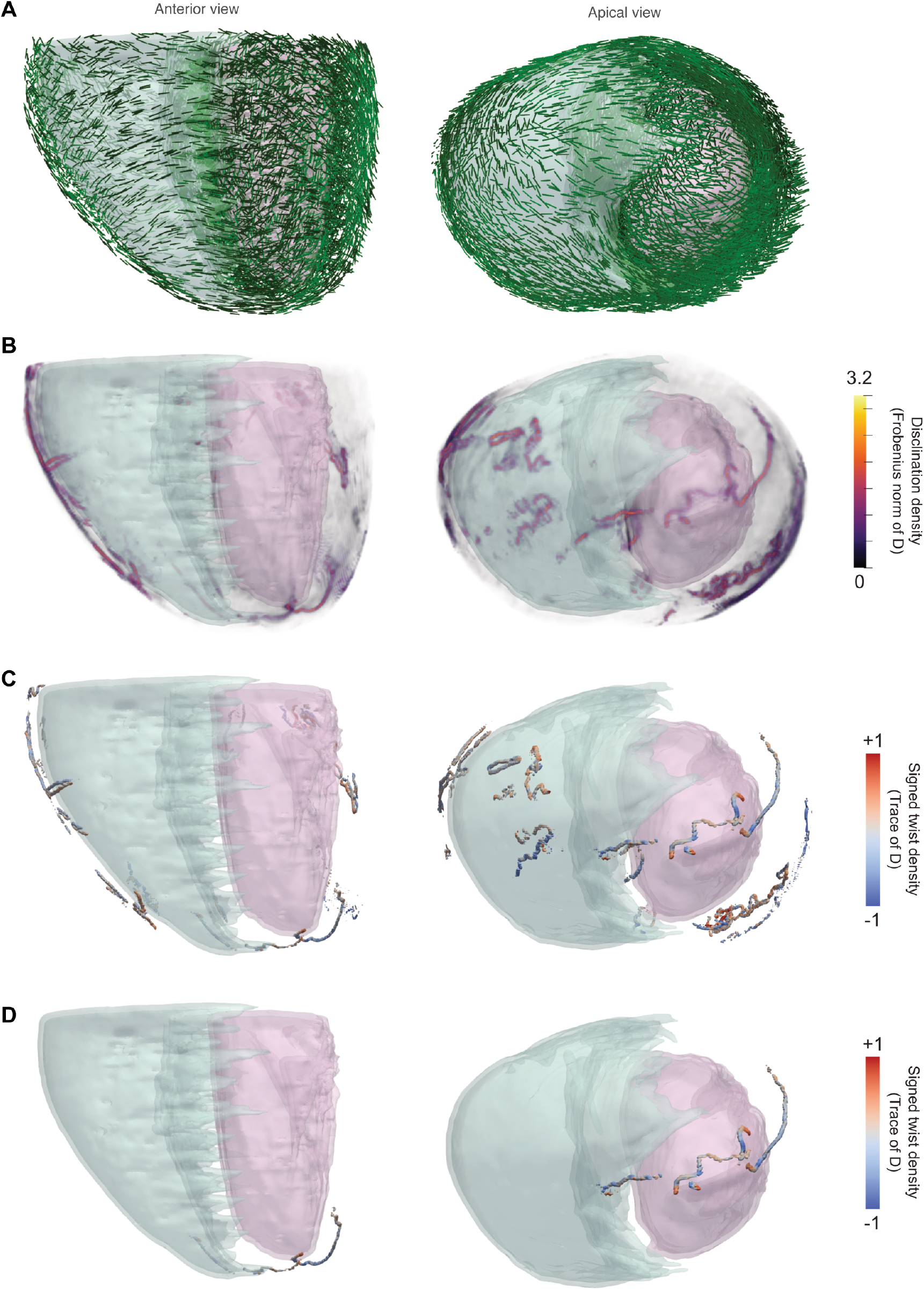
Workflow for the detection of disclination lines. (a) Bar plot representation of the fibre orientation field within the myocardium based on MRI data. Each bar indicates the local fibre direction derived from diffusion tensor imaging. (b-d) Computational pipeline for detecting disclination lines. The cleaning procedure removes artefactual features that do not represent topologically significant disclinations, including: boundary-related fluctuations near tissue surfaces, poorly resolved regions of the right ventricle (RV) where the wall thickness is too thin, and small surface loops that can be trivially resolved. (b) Frobenius norm of the disclination density tensor throughout the myocardium, showing fibre orientational disorder. (c) Extracted cores of regions where the Frobenius norm exhibits significantly large values, identifying potential singular sites. (d) Final result after the cleaning procedure, displaying only the disclination lines that are topologically relevant for cardiac fibre organisation.

**Extended Data Fig. 2.**
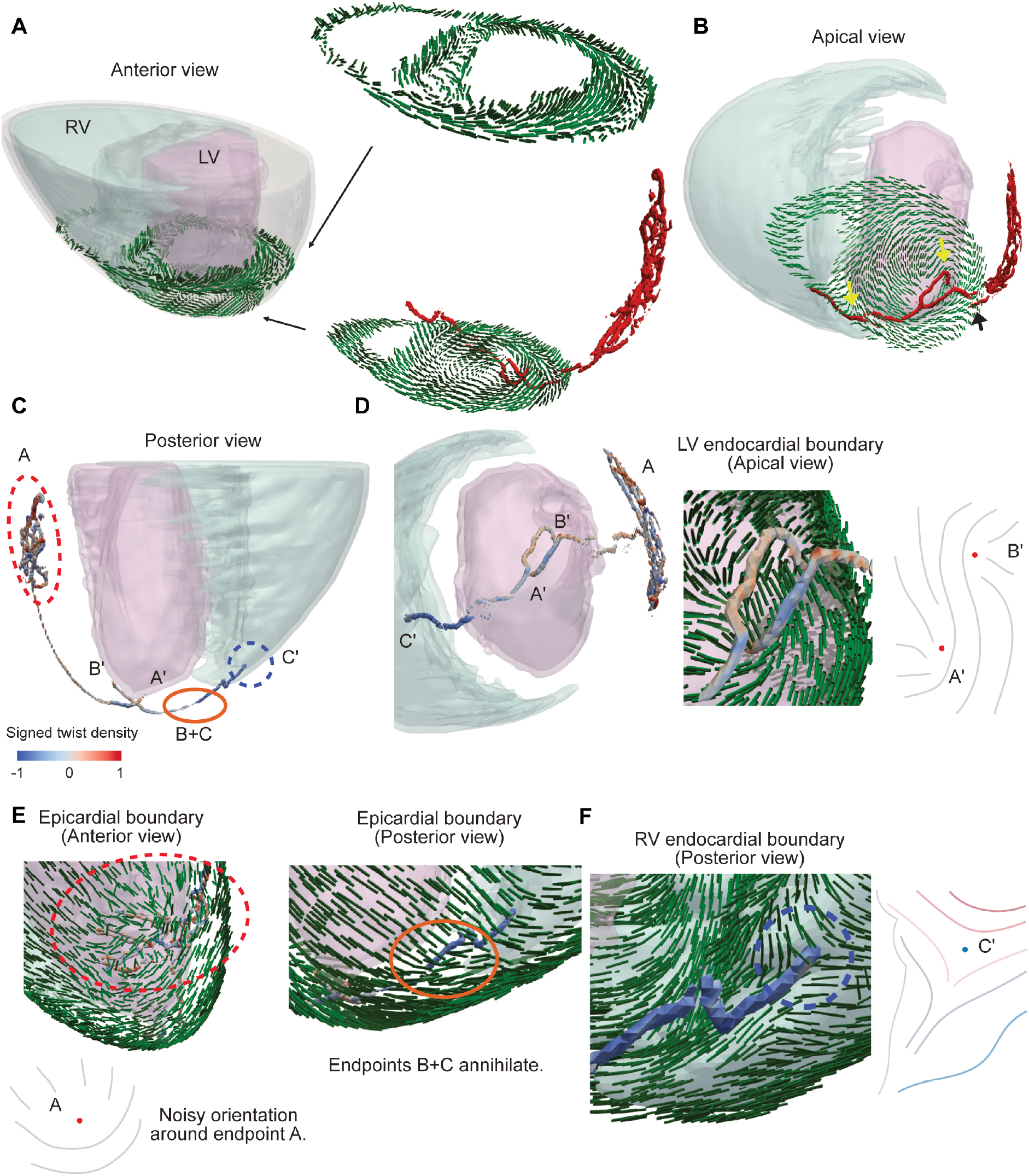
Topological characterisation of three-dimensional fibre architecture in the adult mouse heart (the second sample). (a) Three-dimensional nematic pattern of cardiac fibre structure in the adult mouse heart, showing transmural twist and disclination lines. Transmural twist demonstrates CCW twist. (b) Apical view of a cross-section of the orientation field with disclination lines. Two-dimensional defect-like regions are indicated by arrows. (c) Magnified view of disclination lines from the posterior side. Heat colour represents the signed twist density. (d) Orientation field surrounding disclination line endpoints in the left ventricle. (e) Sub-epicardial orientation pattern in the vicinity of disclination line endpoints. While the orientation field is noisy around the endpoint A in this sample, the total surface charge of A is certainly +1/2. The endpoints B and C observed in the first sample in Fig.1 annihilate and the disclination lines B-B’ and C-C’ are connected. (f) Magnified view of the endocardial endpoint of the disclination line on the right ventricle.

**Extended Data Fig. 3.**
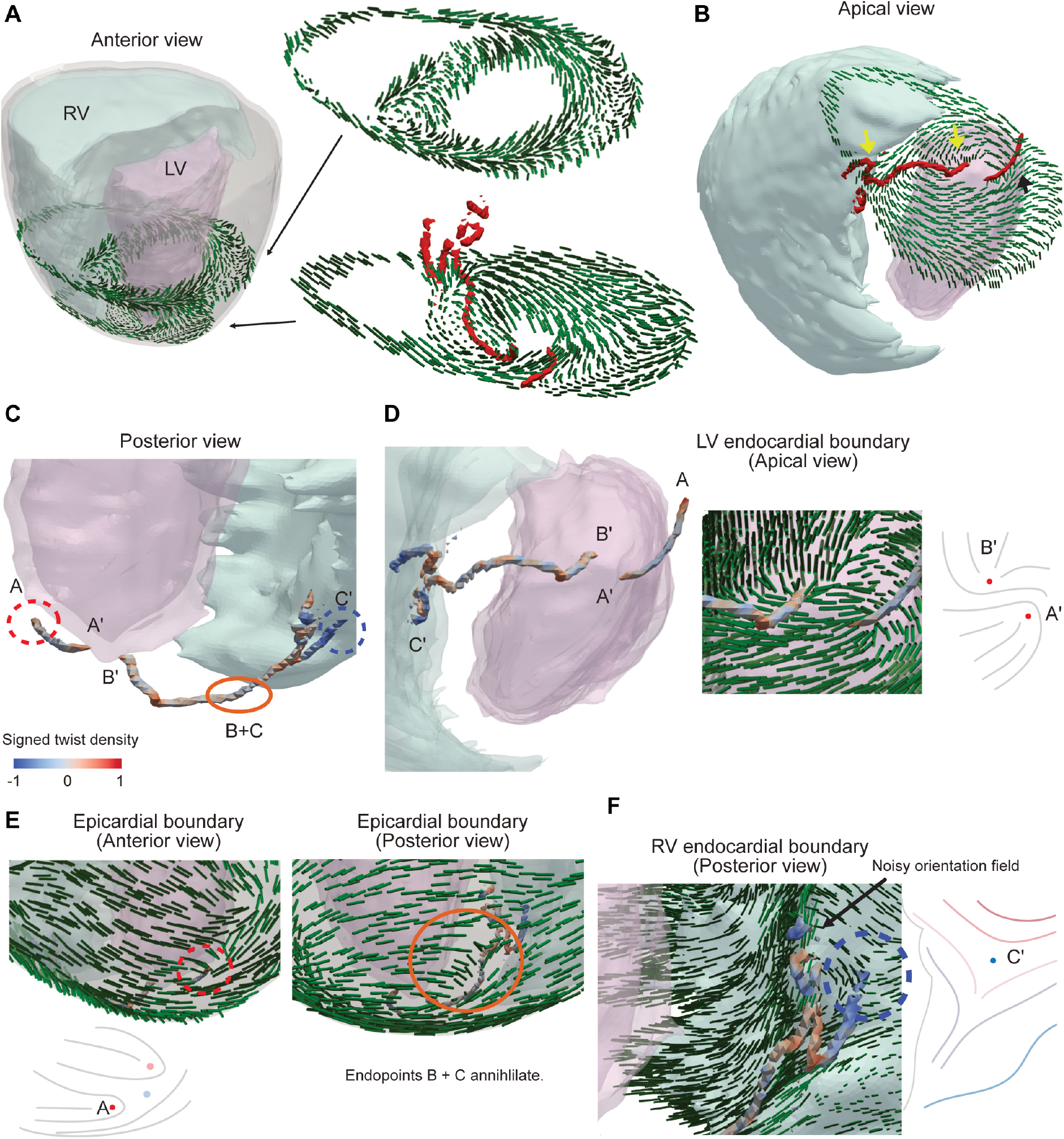
Topological characterisation of three-dimensional fibre architecture in the adult mouse heart (the third sample). (a) Three-dimensional nematic pattern of cardiac fibre structure in the adult mouse heart, showing transmural twist and disclination lines. Transmural twist demonstrates CCW twist. (b) Apical view of a cross-section of the orientation field with disclination lines. Two-dimensional defect-like regions are indicated by arrows. (c) Magnified view of disclination lines from the posterior side. Heat colour represents the signed twist density. (d) Orientation field surrounding disclination line endpoints in the left ventricle. (e) Sub-epicardial orientation pattern in the vicinity of disclination line endpoints. We observe a set of two surface defects with charge +1/2 and −1/2 above the endpoint A due to the noisy orientation field around the endpoint A. Nevertheless, the total surface charge around the endpoint A is +1/2 as those of the other two samples. The endpoints B and C observed in the first sample in Fig.1 annihilate and the disclination lines B-B’ and C-C’ are connected. (f) Magnified view of the endocardial endpoint of the disclination line on the right ventricle.

**Extended Data Fig. 4.**
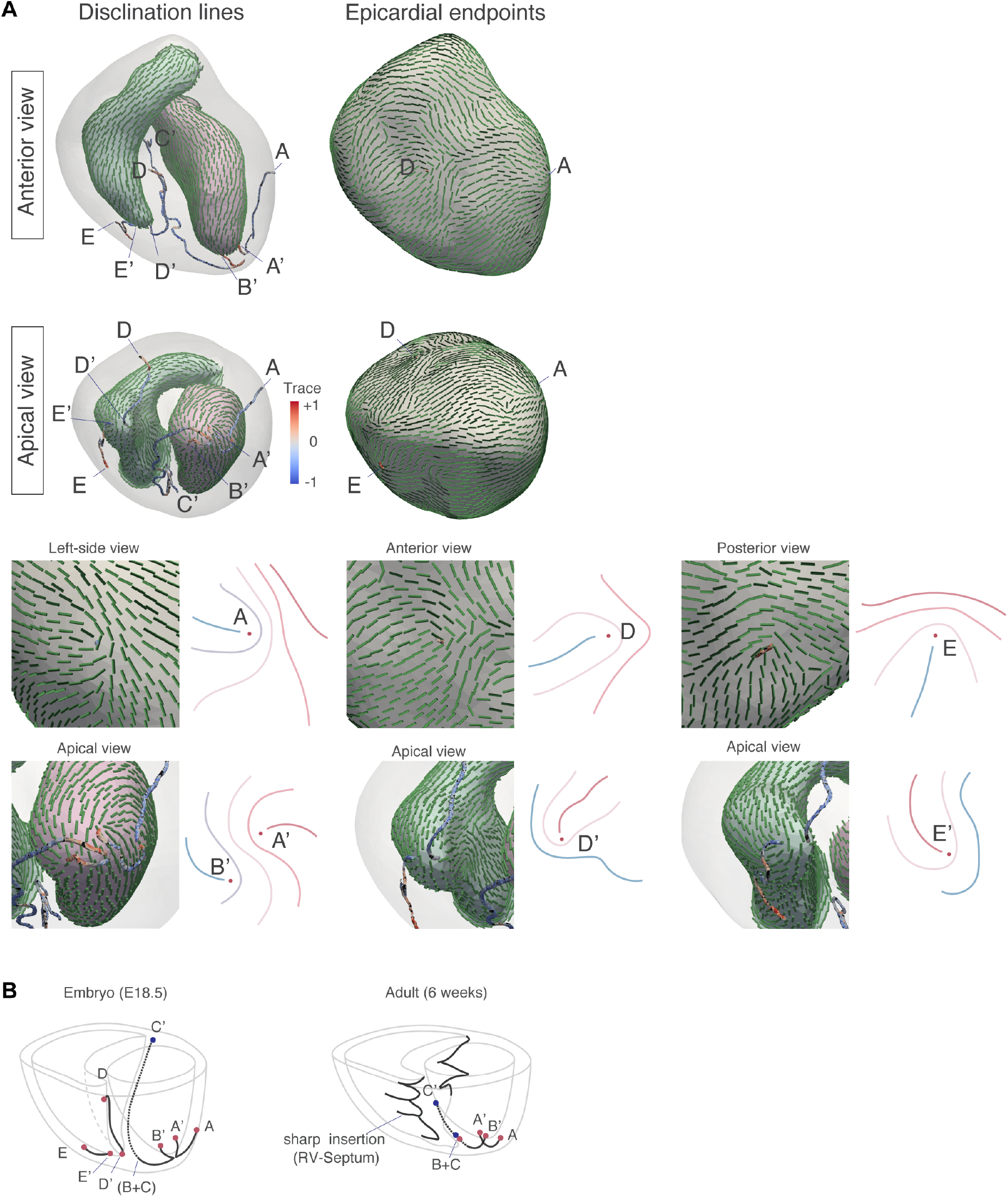
Disclination lines in embryonic heart and conserved topological singularities (a) Disclination line structures in the late-stage embryonic heart (E18.5) are observed at similar positions corresponding to those in the adult heart. At the apex, two disclination lines originate from the apex to the endocardial surface (A’ and B’). In the septum, more distinct disclination lines were identified in the embryo. The relationship between the disclination line running posteriorly through the septum and the apex (B’-C’) exhibits a structural similarity to that observed in adult samples (Fig. S2 and S3). Additionally, disclination lines were identified near the RV (D-D’, E-E’). (b) Comparison of disclination line structures between embryo and adult. In the adult heart, the right ventricle (RV)-interventricular septum (IVS) junction forms a spike-like protrusion, where the acute angular geometry at the front edges imposes boundary conditions analogous to those of topological defects. Therefore, the disclination line near the RV observed in the embryo (D-D’) is absorbed into the RV-septum edge morphology in the adult. Topological singularities are located at the vortex cordis and either within the septum or at the RV-septal junction, depending on the morphology of RV.

**Extended Data Fig. 5.**
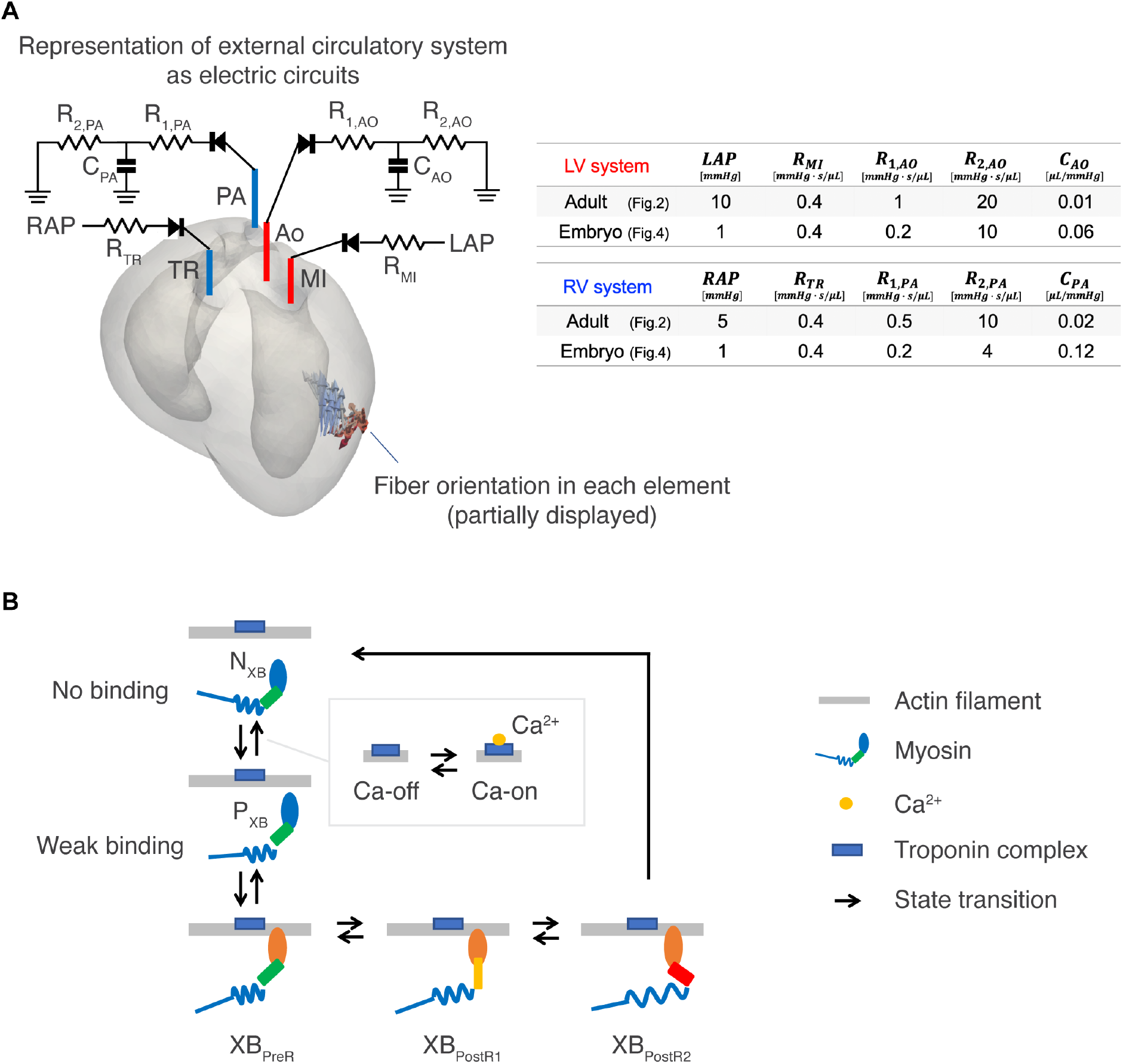
Mechanical modelling and simulation of heart contraction with fibre orientation field. (a) Finite element model of the ventricles and electric circuits representing the circulatory system external to the heart. The ventricular wall is divided into tetrahedral elements, and each element is assigned a vector indicating the fibre direction (arrows in the figure). Preload and afterload were separated so that left ventricular ejection capacity would not be affected by right ventricular ejection capacity. The table shows the parameters for the adult and the embryonic heart simulations. (b) State transition model of the actomyosin system. The small box shows the state transition model of troponin on actin filaments. The state transition rate of troponin depends on intracellular calcium ion concentration, and the state of troponin affects the transition rate between non-attached (N_XB_) and weakly-attached (P_XB_) states of the myosin molecules beneath it. Myosin molecules bound to actin filaments generate force through a two-step lever arm rotation (first step: XB_PreR_ to XB_PostR1_, second step: XB_PostR1_ to XB_PostR2_). Approximately 1000 myosin molecules are placed in each individual finite element constituting the ventricle, and the contractile force acting in the fibre direction is determined from the resultant force of each myosin molecule in the bound state. The sliding motion of actin filaments is calculated from the contraction and extension motion in the fibre direction, and this is reflected in changes in the mechanical load on each myosin molecule. These changes in mechanical load affect the state transition rates.

**Extended Data Fig. 6.**
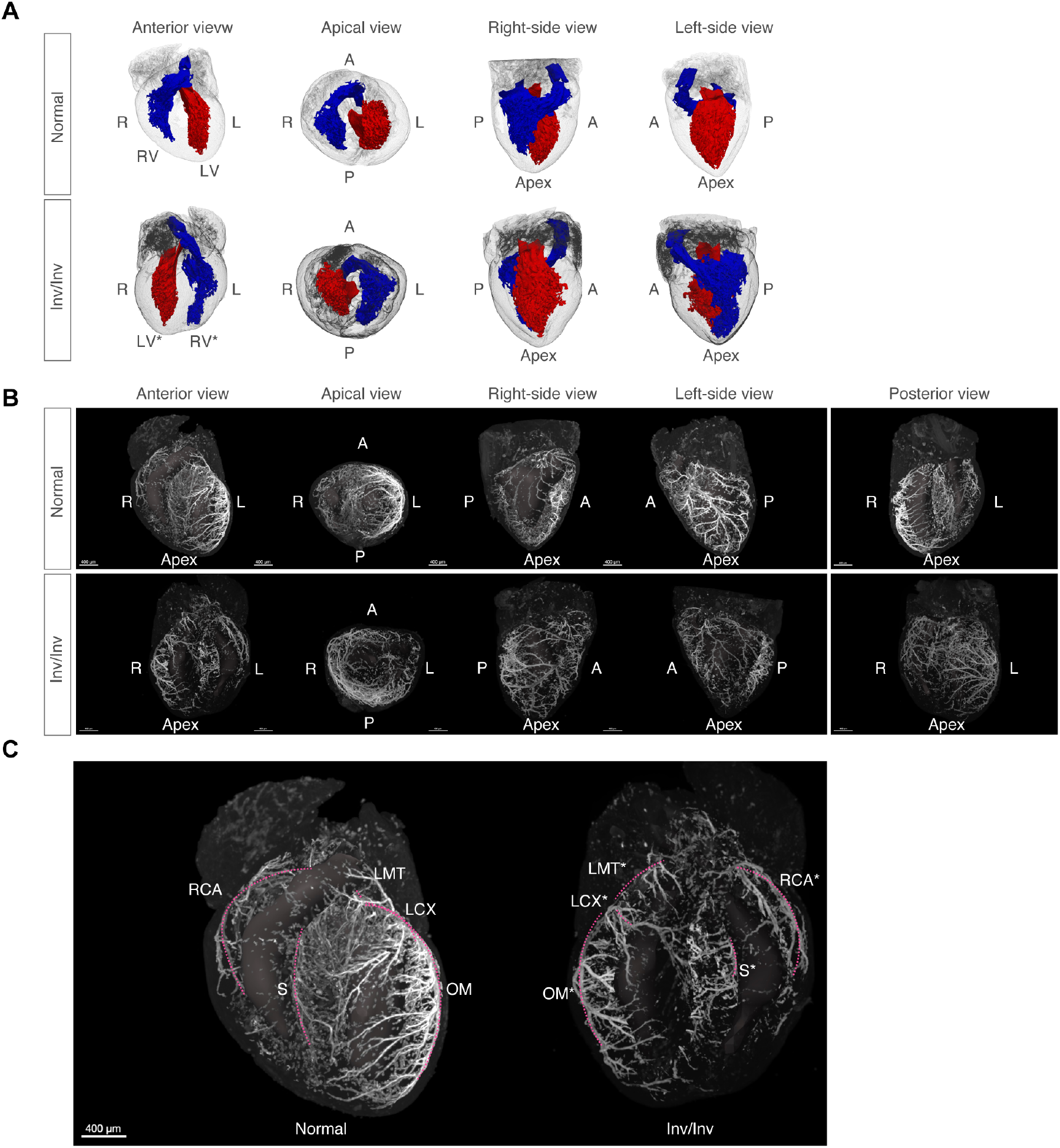
Inverted macroscopic structures in the heterotaxy heart. (a) Ventricles of wild-type and inverted heart. LV* and RV* represent the anatomical left ventricle and anatomical right ventricle in a heterotaxy heart. (b) Coronary arteries of wild-type and inverted heart. Left-right inverted corresponding vessels are present in the heterotaxy heart. (c) Coronary artery annotations. Asterisks indicate corresponding vessels in the heterotaxy heart. LMT: left main trunk, LCX: left circumflex artery, OM: obtuse marginal artery, S: septal artery, RCA: right coronary artery.

**Extended Data Fig. 7.**
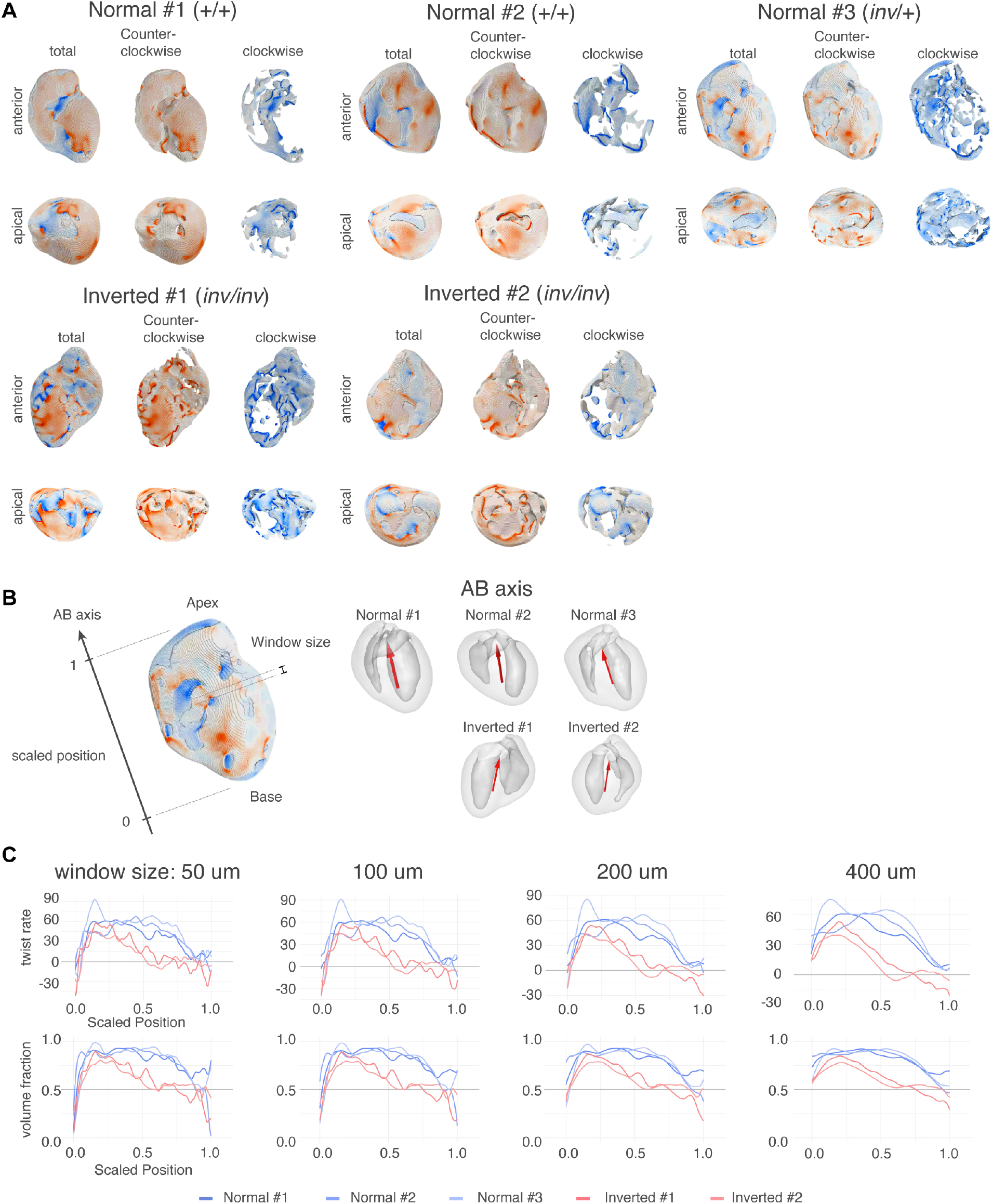
Analysis of twist chirality in normal and inverted hearts. (a) Twist chirality of all normal and inverted hearts. (b) The chirality profile was measured along the apex-to-base orientation. This orientation was determined as the principal axis of the left ventricular shape in each heart. (c) The effect of window size on the chirality profiles. In all cases, the normal heart showed consistent CCW chirality, while inverted hearts showed mixed chirality near the base region.

**Extended Data Fig. 8.**
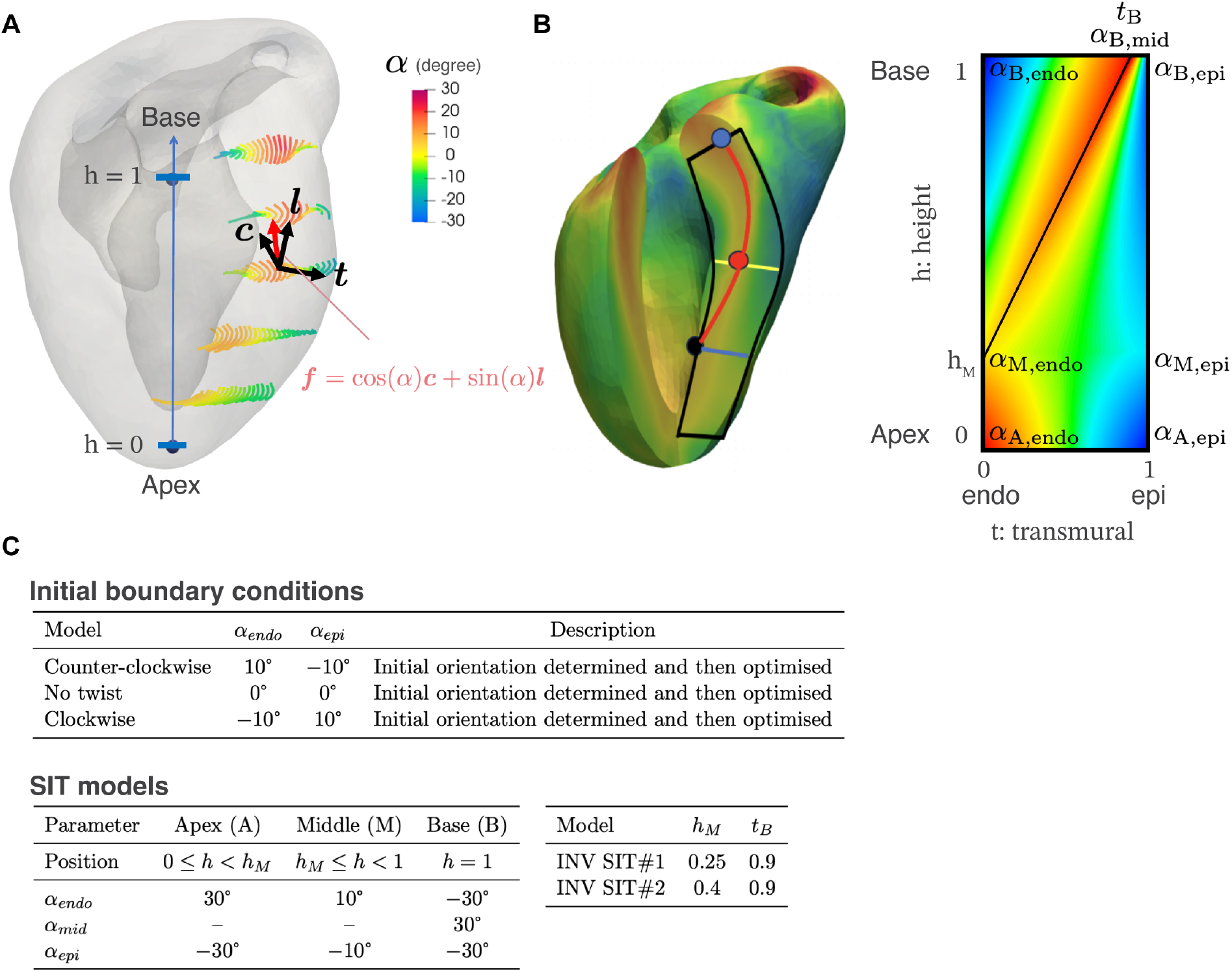
Generation of cardiac fibre structures with artificially controlled chirality for FEM. The shape of an inverted heart is shown here. (a) Fibre angle α was introduced as a function of the height *h* and the transmural position *t*. Here, ***f*** is the unit vector of the orientation field, ***c*** is the unit vector in the circumferential direction, and ***l*** is the unit vector in the longitudinal direction. The initial orientation was determined by specifying α at the endocardium and epicardium as boundary conditions, followed by linear interpolation in the transmural direction. (b) For the SIT model, additional control points α_M,endo_, α_B,mid_ were introduced. First, linear interpolation was performed between α_M,endo_ at (*h, t*) = (*h*_*M*_, 0) and α_B,mid_ at (*h, t*) = (1, *t*_*B*_). Subsequently, at each height *h*, the fibre angle within the wall was generated by linear interpolation in the transmural direction according to α_endo_ and α_epi_. (c) Table of parameters used for the mouse embryonic hearts. The optimisation process was performed to increase the impulse following the previous studies. It was conducted with weak contractile force (4 kPa) adapted for the embryonic mouse. Through this optimisation, the initial slight twist tended to increase, generating the final orientation field.

**Extended Data Fig. 9.**
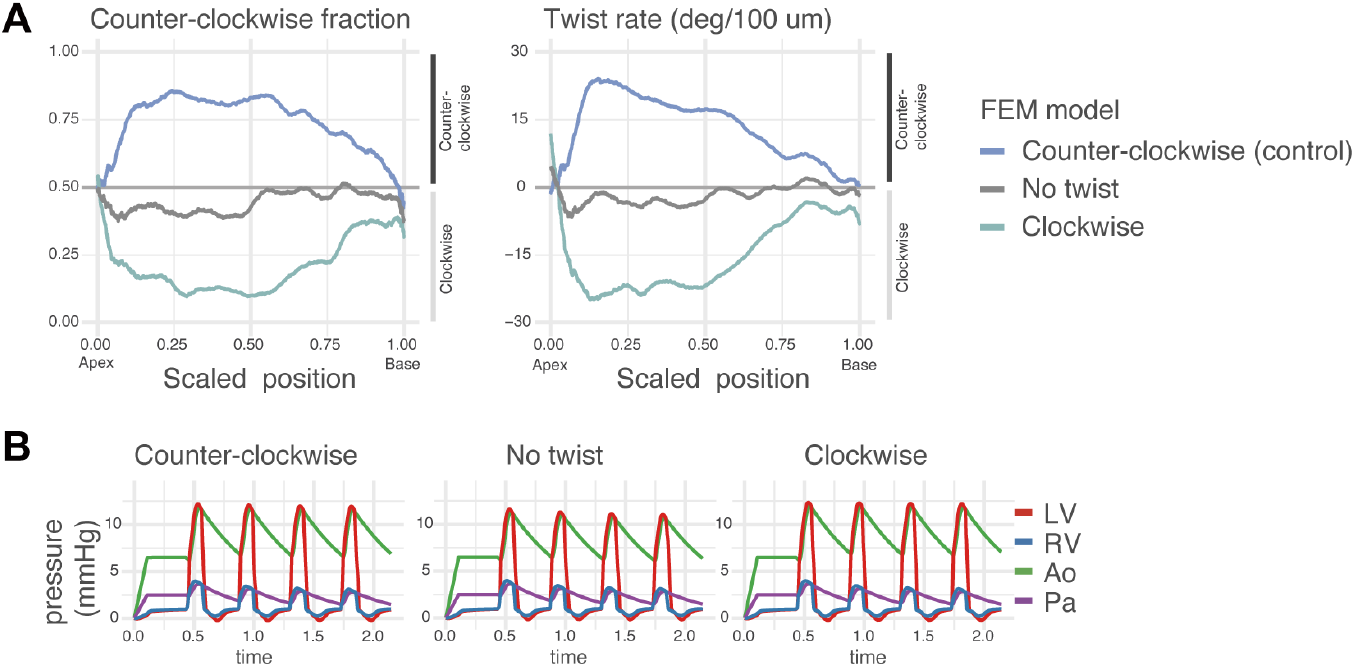
(a) Profiles of twist chirality of the fibre structures in FEM models in Fig. 4a. In addition to a similar profile to the normal sample in Fig. 3g, inverted chirality and minimal chirality are observed. These structures are generated from different initial boundary conditions as described in Fig. S8. (b) Time course of blood pressures during beating simulation. LV, left ventricle; RV, right ventricle; Ao, aorta; Pa, pulmonary artery.

**Extended Data Fig. 10.**
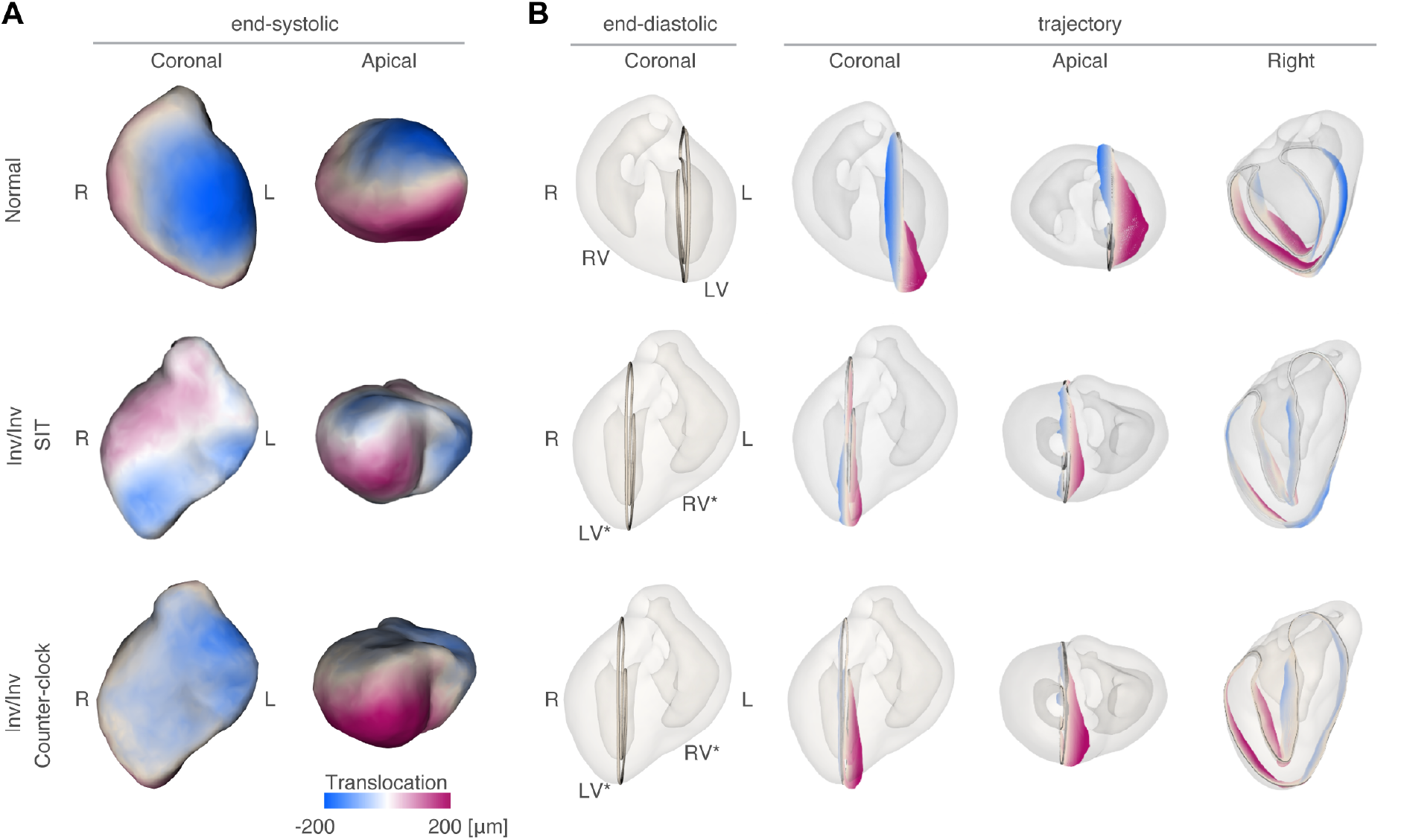
Cardiac motion during contraction for WT heart, situs inversus totalis heart and inverted heart with artificially uniform CCW twist. Displacement during contractile motion was represented with reference to the position in the end-diastolic heart for each tissue, with the base of the heart fixed in position. (a) Magnitude and direction of motion of the superficial tissue. (b) Traced motion of the sagittal section of the left ventricle (LV) and the anatomical left ventricle (LV*). The normal and *inv*/*inv* CCW hearts possess consistent CCW chirality and exhibit twisting motion of the entire organ in the same direction, despite the left-right inversion of the organ shape. In contrast, in SIT, while the motion of the apex is similar to that in other conditions, a reversed motion is observed in the basal region. See also Extended Data Movie 1.

